# Navigome: Navigating the Human Phenome

**DOI:** 10.1101/449207

**Authors:** Héléna A Gaspar, Christopher Hübel, Jonathan R. I. Coleman, Ken Hanscombe, Gerome Breen

## Abstract

We now have access to a sufficient number of genome-wide association studies (GWAS) to cluster phenotypes into genetic-informed categories and to navigate the “phenome” space of human traits. Using a collection of 465 GWAS, we generated genetic correlations, pathways, gene-wise and tissue-wise associations using MAGMA and S-PrediXcan for 465 human traits. Testing 7267 biological pathways, we found that only 898 were significantly associated with any trait. Similarly, out of ~20,000 tested protein-coding genes, 12,311 genes exhibited an association. Based on the genetic correlations between all traits, we constructed a phenome map using t-distributed stochastic neighbor embedding (t-SNE), where each of the 465 traits can be visualized as an individual point. This map reveals well-defined clusters of traits such as education/high longevity, lower longevity, height, body composition, and depression/anxiety/neuroticism. These clusters are enriched in specific groups of pathways, such as lipid pathways in the lower longevity cluster, and neuronal pathways for body composition or education clusters. The map and all other analyses are available in the Navigome web interface (https://phenviz.navigome.com).

## Introduction

Genome-wide association studies (GWAS) test associations between common genetic variants and phenotypes, such as metabolic disorders, psychiatric disorders^1^, or personality traits. GWAS summary statistics summarize genome-wide associations between a phenotype and common variants - mainly single nucleotide polymorphisms (SNPs). Unlike individual genotypes, GWAS summary statistics are often freely accessible online. Hundreds of GWAS summary statistics are now publicly available, forming a genetic-informed phenotypic space or phenome.

In recent years, methods have been designed to harness GWAS summary statistics to estimate the genetic correlation between traits, as well as their shared biological pathways and shared genes. Linkage disequilibrium score regression (LDSC)^2^ can estimate the genetic correlation between two traits by relying on the fact that SNPs in high linkage disequilibrium (LD) have, on average, stronger associations than SNPs in low LD. In 2015, an atlas of genetic correlations for 24 traits was published by Bulik-Sullivan et al^3^; in 2016, an online tool (LD Hub^4^) provided a way to easily compute genetic correlations between a GWAS and thousands of others. To understand which biological pathways may be implicated in each trait investigated in a GWAS, pathway analysis methods accounting for LD have been developed and are implemented in softwares such as INRICH^5^ or MAGMA^6^. These approaches calculate the overall association of a set of genes (i.e., pathway) with a phenotype, thereby providing insight into potential biological mechanisms. At the gene level, genetic associations can be inferred by combining SNP p-values derived from GWAS, while accounting for LD. The tissue-specific expression of those genes can be predicted by exploiting expression quantitative loci (eQTLs) with software such as S-PrediXcan^7^.

These four layers of analyses (correlations, pathways, genes, and their expression) can be used to identify and understand biological mechanisms associated with each phenotype. To visualize the correlation structure of the human phenome, we propose the use of dimensionality reduction method t-distributed stochastic neighbor embedding^8^ (t-SNE). A t-SNE phenome map can be used to define clusters of phenotypes based on genetic information, and the genetic characteristics of these clusters can be investigated by exploring their (shared) biological pathways and gene associations. GWAS studies have now reached a number to make it possible to find phenotype clusters, and better understand the relationship between them. To allow easy browsing of this data, we developed the Navigome web interface (http://phenviz.navigome.com) associated in this paper, gathering a phenome map, genetic correlations, visualizations of significant pathways, significant genes and gene-tissue pairs.

## Results

### Phenome map and phenotype clusters

Seven large and well-defined clusters could be identified from the t-SNE phenome map constructed from genetic correlations (Figure 1): (1) body composition, (2) height and related measurements, (3) depression, anxiety and neuroticism, (4) phenotypes including years of education and increased longevity (longevity +), (5) lower longevity (longevity -), (6) metabolites in HDL particles, (7) metabolites in VLDL particles, and (8) metabolites in IDL, LDL, and some metabolites in very small VLDL particles. The phenotypes contained in these clusters can be found in Supplementary Figures 1-8. These clusters could still be observed when varying the t-SNE perplexity parameter (cf. https://www.phenviz.navigome.com/map). Clusters 3 (depression and anxiety) and 5 (lower longevity) could arguably be grouped together; cluster 5 includes ADHD, coronary artery disease and social deprivation, and could be seen as the negative counterpart of the education and longevity (+) cluster. Interestingly, metabolites in different lipoprotein particles (HDL, VLDL, IDL and LDL) are clustered by type of lipoprotein rather than by metabolite category. Among phenotypes separate from these large and coherent clusters, schizophrenia and bipolar disorder are mapped in the same small cluster, which has a variable position on the map when changing the perplexity parameter - it is however mapped between the depression and anxiety cluster and the education cluster.

**Figure 1.**
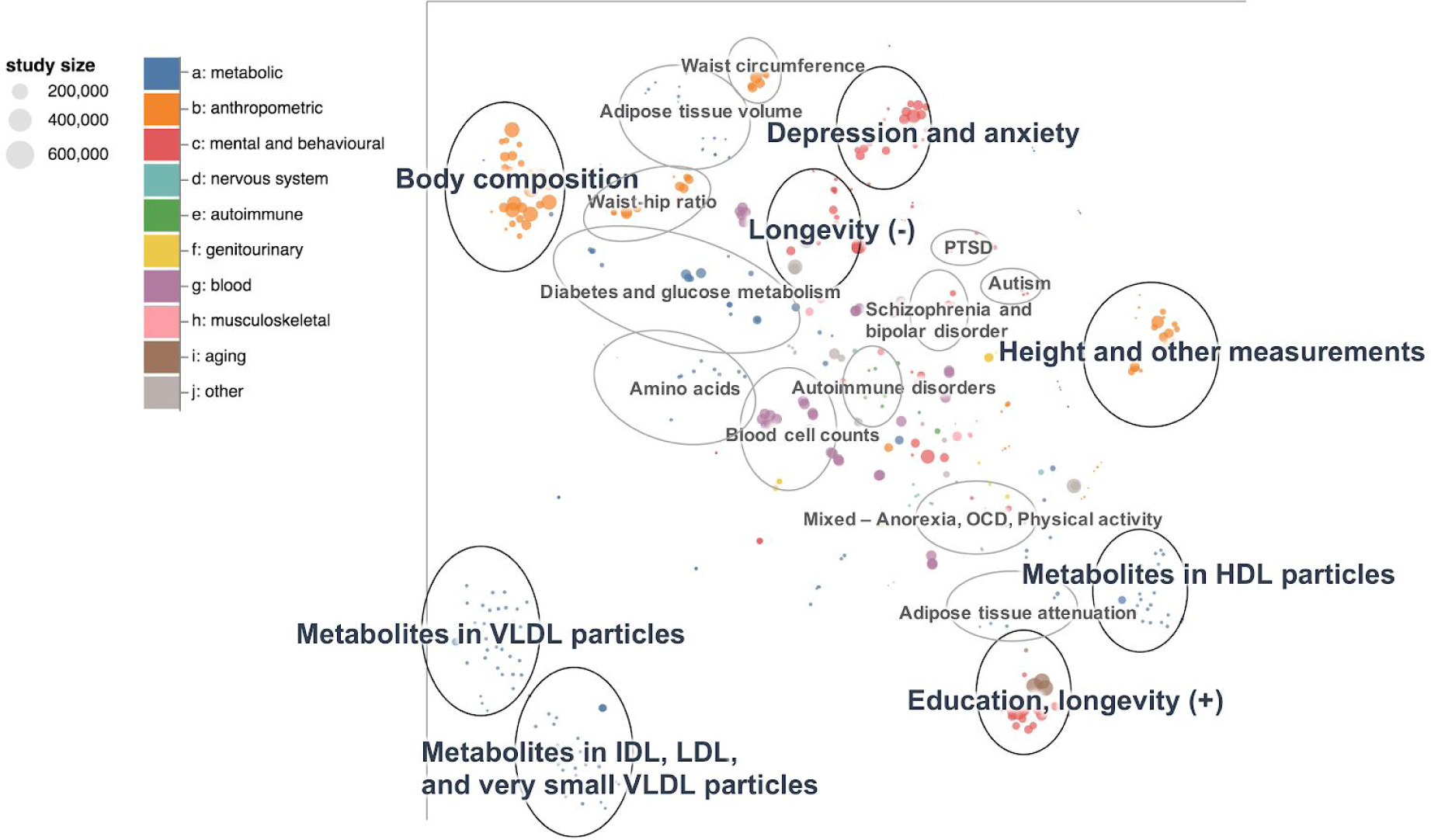
Phenome map of genetic correlations. The map was constructed using the t-distributed stochastic neighbor embedding algorithm (t-SNE) with perplexity = 30, based on the genetic correlation matrix of 465 traits, computed using genome-wide association study (GWAS) statistics. Large coherent clusters are highlighted in black, whereas some other clusters of interest are highlighted in grey. The colours correspond to the trait categories, and the marker size to the GWAS sample size.

By colouring phenotypes by correlation with years of education (Figure 2A), the map becomes clearly separated into two distinct parts: one that is positively correlated with phenotypes related to education and longevity, and the other negatively. Among negatively correlated phenotypes, we can find the depression and anxiety cluster; among positively correlated phenotypes, metabolites in HDL particles are the closest to the education cluster (Figure 2B). Total HDL cholesterol has a correlation pattern across phenotypes similar to years of education (Figure 2B), but differs in that it shows positive rather than negative correlation in the depression and anxiety cluster.

**Figure 2.**
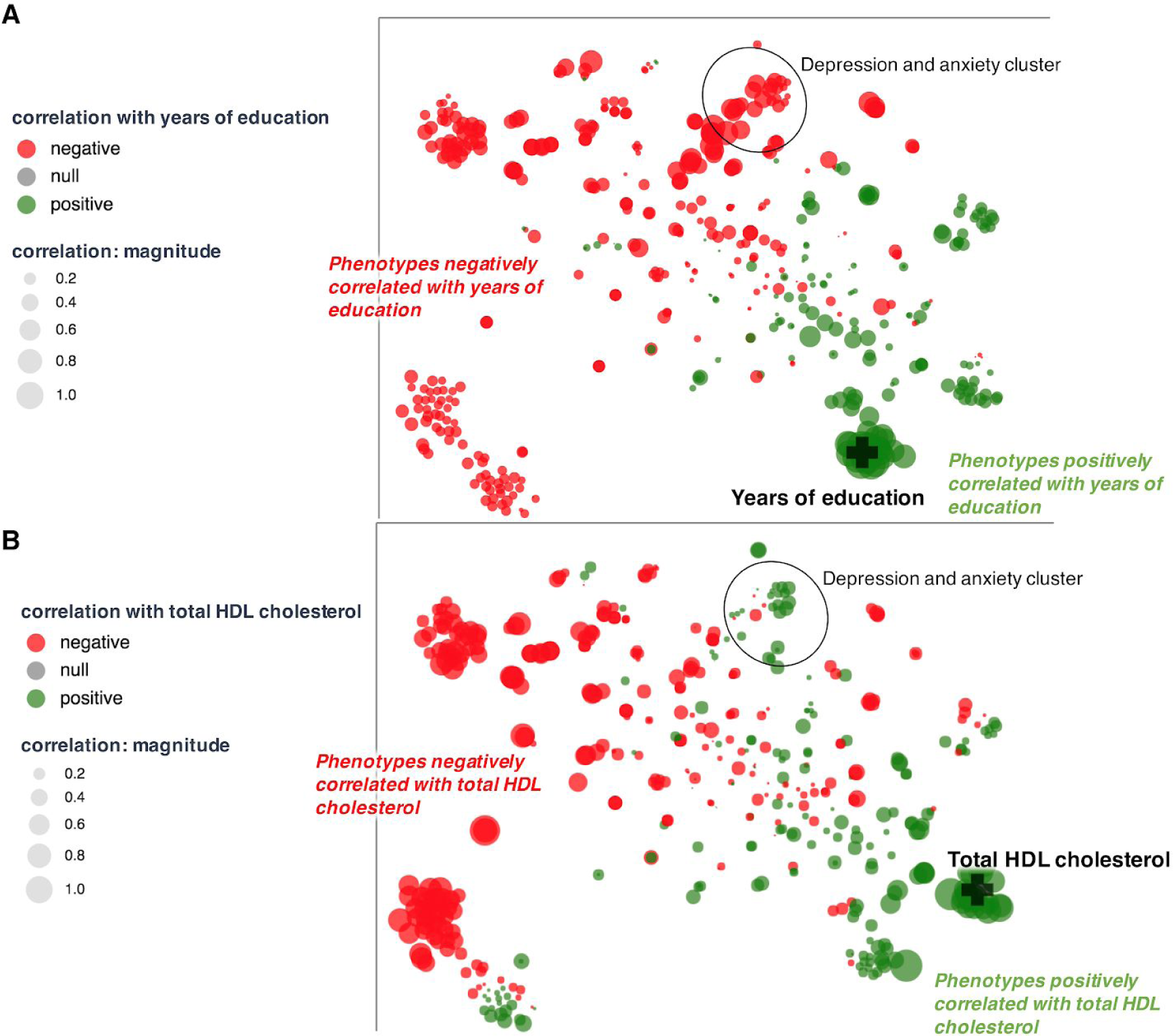
Phenome map of genetic correlations (cf. Figure 1) coloured by (A) correlation with years of education, and (B) correlation with HDL cholesterol.

Phenotypes coloured by correlation with schizophrenia (Figure 3) show a more complex pattern on the map - correlations in the depression and anxiety cluster are strongly positive, but correlations in the education and longevity cluster are mixed: schizophrenia is negatively correlated with intelligence and longevity but positively correlated with years of education.

**Figure 3.**
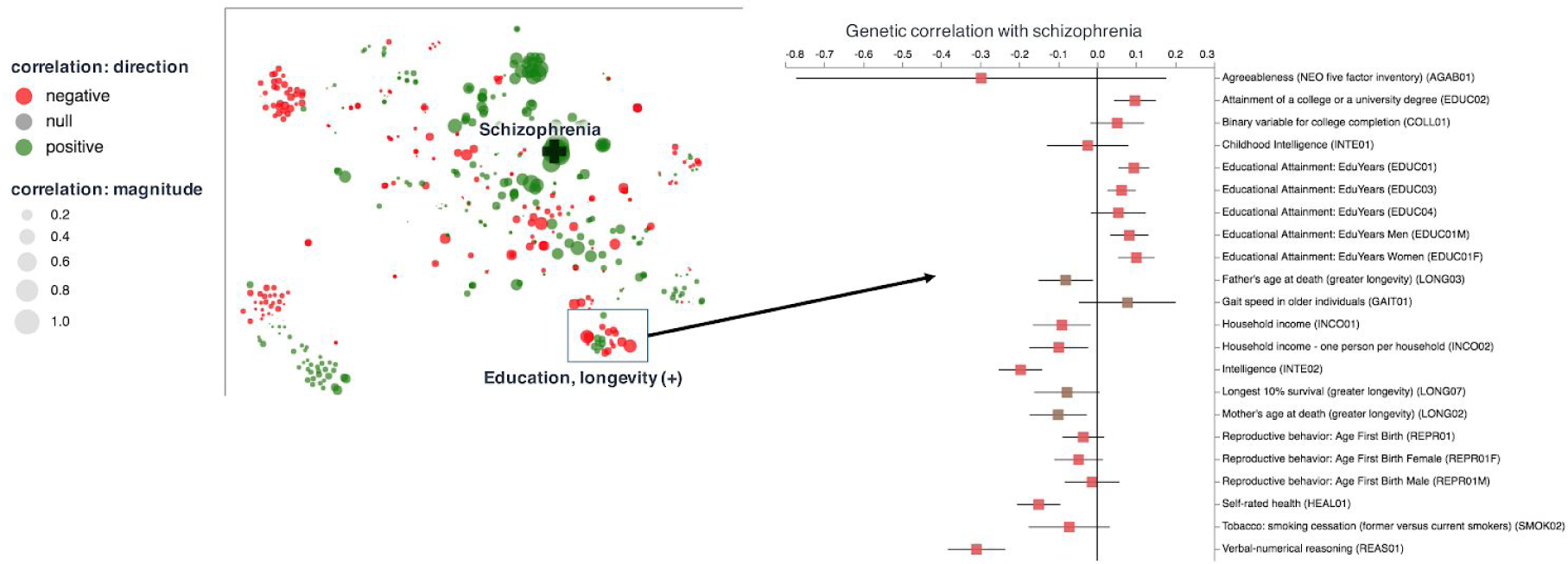
Phenome map of genetic correlations (cf. Figure 1) coloured by correlation with schizophrenia (PGC2), with highlighted genetic correlations in the education/longevity cluster.

### Gene and pathway associations

There are 99,378 gene-phenotype associations with p < 2.5E-6 for all the phenotypes in at least one MAGMA or S-PrediXcan analysis, corresponding to 12,311 unique genes. MAGMA alone identifies 84,856 associations for 11,078 genes (p ≤ 2.5E-6), whereas S-PrediXcan identifies 51,375 associations for 8087 distinct genes (p ≤ 2.5E-6), or 36,647 associations for 6589 distinct genes with a more stringent p-value threshold taking into account all gene-tissue pairs (p ≤ 1.9342E-7). This means that at the same significant threshold, S-PrediXcan identifies 14,522 gene-phenotype associations not identified by MAGMA. Visualizations of MAGMA and S-PrediXcan results and their overlap can be found in Navigome (phenviz.navigome.com); a “gene profile” can be generated to visualize S-PrediXcan results across phenotypes.

3460 phenotype-pathway associations are significant (p ≤ 1.0E-5), corresponding to 898 discrete unique pathways. Pathways occurring at least two times in education/longevity, depression and anxiety, metabolites in HDL, body composition and height clusters are reported in Tables 1-3.

**Table 1:**
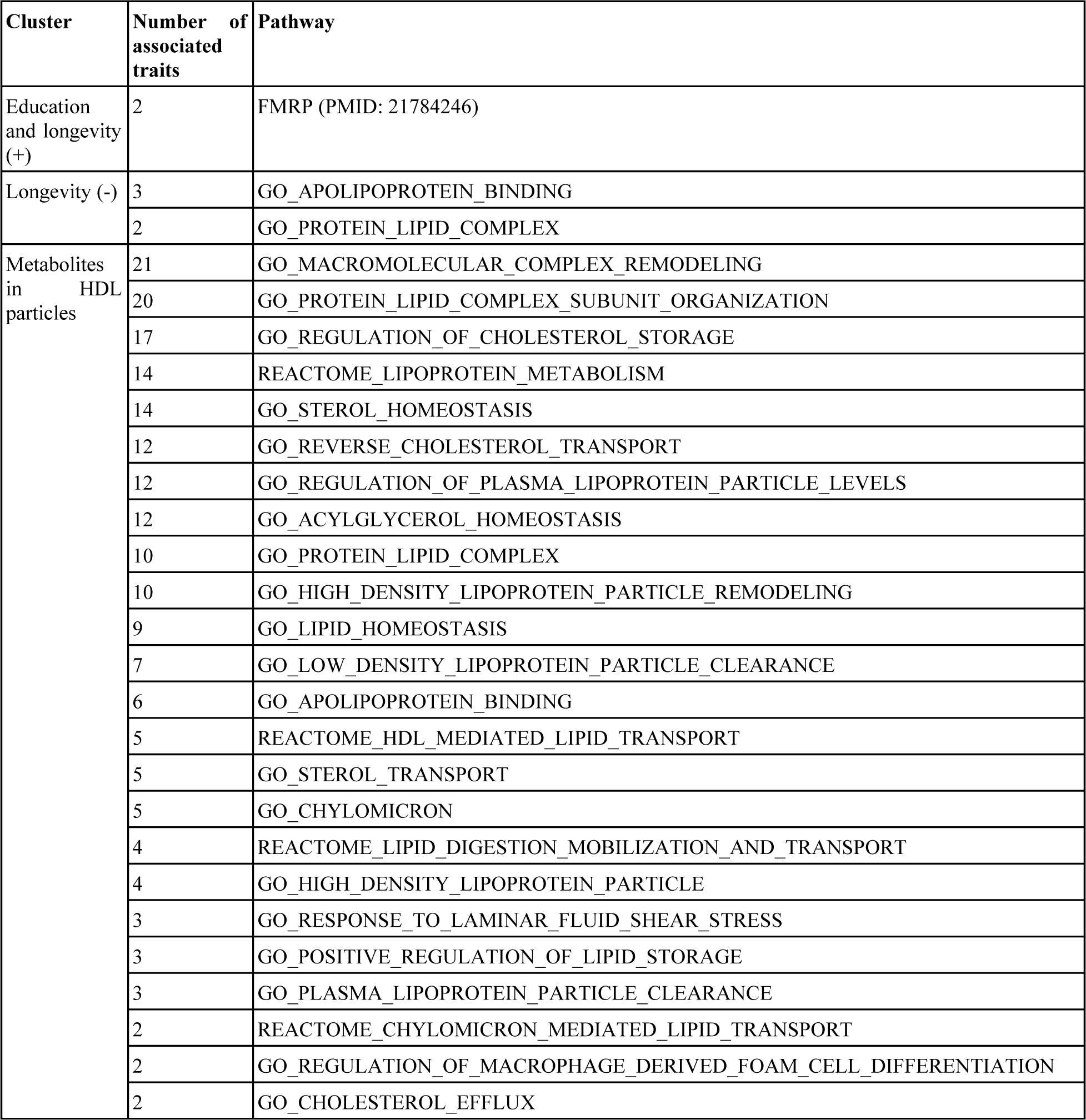
Most frequent pathways in education, longevity and HDL clusters on the phenome map.

**Table 2:**
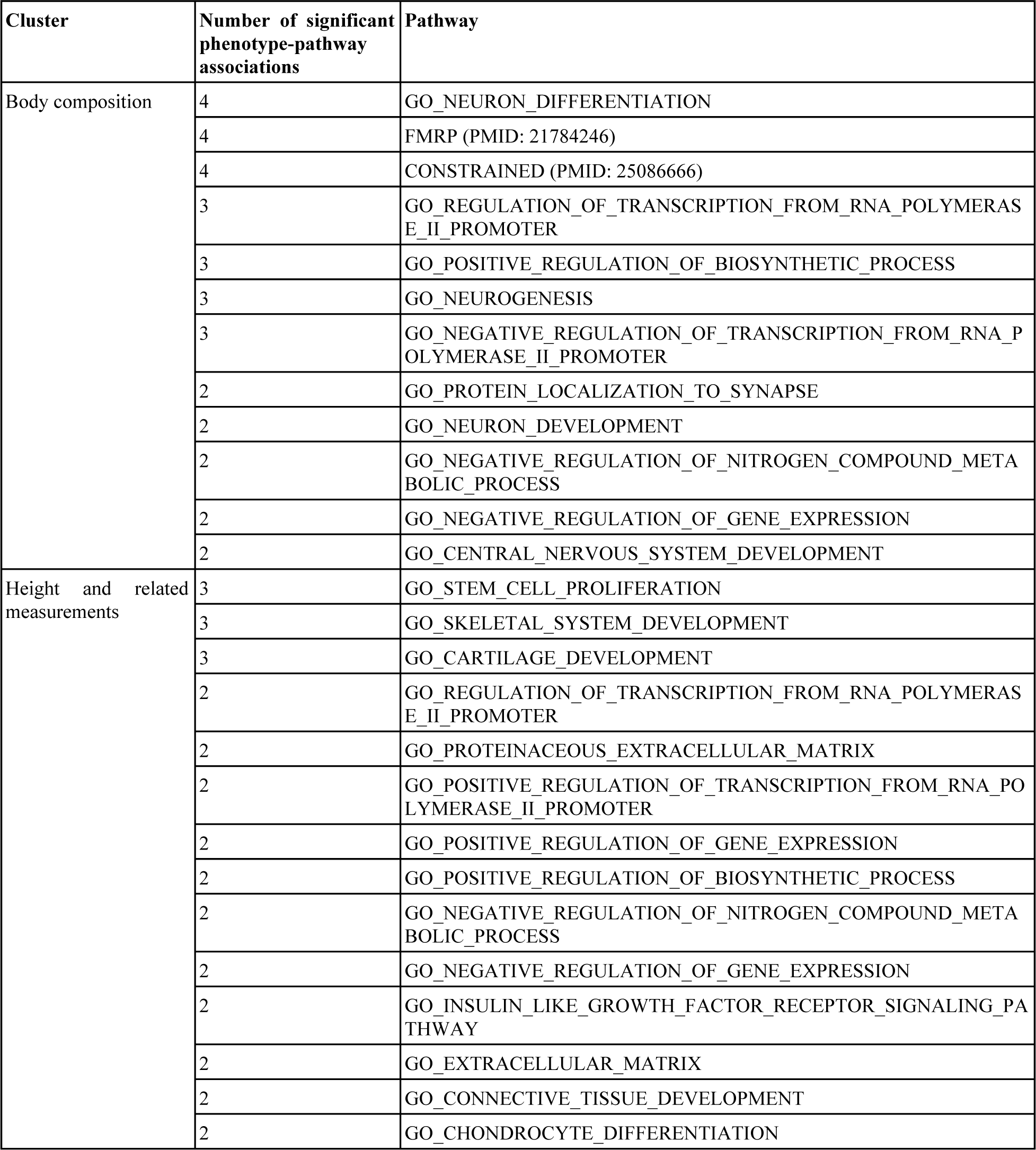
Most frequent pathways in body composition and height cluster on the phenome map, and the n

| Cluster | Number of significant phenotype-pathway associations | Pathway |
| --- | --- | --- |
| Body composition | 4 | GO_NEURON_DIFFERENTIATION |
|  | 4 | FMRP (PMID: 21784246) |
|  | 4 | CONSTRAINED (PMID: 25086666) |
|  | 3 | GO_REGULATION_OF_TRANSCRIPTION_FROM_RNA_POLYMERASE_II_PROMOTER |
|  | 3 | GO_POSITIVE_REGULATION_OF_BIOSYNTHETIC_PROCESS |
|  | 3 | GO_NEUROGENESIS |
|  | 3 | GO_NEGATIVE_REGULATION_OF_TRANSCRIPTION_FROM_RNA_POLYMERASE_II_PROMOTER |
|  | 2 | GO_PROTEIN_LOCALIZATION_TO_SYNAPSE |
|  | 2 | GO_NEURON_DEVELOPMENT |
|  | 2 | GO_NEGATIVE_REGULATION_OF_NITROGEN_COMPOUND_METABOLIC_PROCESS |
|  | 2 | GO_NEGATIVE_REGULATION_OF_GENE_EXPRESSION |
|  | 2 | GO_CENTRAL_NERVOUS_SYSTEM_DEVELOPMENT |
| Height and related measurements | 3 | GO_STEM_CELL_PROLIFERATION |
|  | 3 | GO_SKELETAL_SYSTEM_DEVELOPMENT |
|  | 3 | GO_CARTILAGE_DEVELOPMENT |
|  | 2 | GO_REGULATION_OF_TRANSCRIPTION_FROM_RNA_POLYMERASE_II_PROMOTER |
|  | 2 | GO_PROTEINACEOUS_EXTRACELLULAR_MATRIX |
|  | 2 | GO_POSITIVE_REGULATION_OF_TRANSCRIPTION_FROM_RNA_POLYMERASE_II_PROMOTER |
|  | 2 | GO_POSITIVE_REGULATION_OF_GENE_EXPRESSION |
|  | 2 | GO_POSITIVE_REGULATION_OF_BIOSYNTHETIC_PROCESS |
|  | 2 | GO_NEGATIVE_REGULATION_OF_NITROGEN_COMPOUND_METABOLIC_PROCESS |
|  | 2 | GO_NEGATIVE_REGULATION_OF_GENE_EXPRESSION |
|  | 2 | GO_INSULIN_LIKE_GROWTH_FACTOR_RECEPTOR_SIGNALING_PATHWAY |
|  | 2 | GO_EXTRACELLULAR_MATRIX |
|  | 2 | GO_CONNECTIVE_TISSUE_DEVELOPMENT |
|  | 2 | GO_CHONDROCYTE_DIFFERENTIATION |

## Discussion

The phenome map can be separated into two parts: one that is more correlated with higher education, greater longevity and height, and another that is more correlated with increased BMI, lower longevity and metabolic or mental disorders. However, there are some caveats. First, a single t-SNE map is not sufficient to make assumptions about the topology of the phenome. This is why we constructed t-SNE maps with different values of the t-SNE perplexity parameter, which is analogous to a “number of neighbours” and can tune the local-global topology balance. The clusters described in the results section were found for all perplexity values. This is an important feature of cluster analysis - several visualizations of a multi-dimensional space provide a better picture of the data structure.

Moreover, the phenome map was built using signed genetic correlations - which means that the phenotype encoding should be taken into account when interpreting results. Inverting alleles for each genetic marker would give a genetic correlation with the same magnitude but opposite sign. Two phenotypes that are the same but with inverted genetic correlations will be mapped far from each other on the map. For example, some longevity GWASs correspond to greater longevity or lower longevity - they are referred on the figures as longevity (+) or longevity (-). Phenotypes in the education and longevity (+) cluster are correlated with better education and greater longevity, whereas phenotypes in the longevity (-) cluster are correlated with lower longevity. Interestingly, the most frequent significant pathways in the longevity (-) cluster are metabolic (*GO_APOLIPOPROTEIN_BINDING* and *GO_PROTEIN_LIPID_COMPLEX*), whereas the most frequent significant pathway in the longevity (+) cluster is synaptic (*FMRP*).

Another caveat is that most of these genetic correlations are not significant - they might have a high magnitude but also high standard error. This is, however, also the advantage of using the phenome map, which allows users to check if phenotypes in a given cluster all have the same correlation direction, even though those correlations may not be significant. The phenome map can also help to differentiate phenotypes which are very close genetically and would be expected to have similar symptoms (such as depression and anxiety) from phenotypes which have some features in common but display different correlations patterns with some phenotype categories. For example, HDL cholesterol and educational years have generally similar correlation patterns; however, they show opposite correlations with phenotypes in the depression and anxiety cluster. Although the HDL and education cluster are neighbours on the phenome map, their biological pathways may be quite different - we find metabolic pathways in the HDL cluster and synaptic pathways in the education cluster.

This type of map can be used to categorize phenotypes genetically, and find pathways and genes which are characteristic of these new phenotype clusters. For example, without prior knowledge on the genetic structure, we categorized body composition and height as “anthropometric traits”. The body composition and height cluster are, however, very different clusters, each characterized by different biological pathways: neuronal pathways for body composition, and gene expression, cartilage, and skeletal pathways for height. For some complex disorders search as schizophrenia, explaining correlation patterns might constitute a challenge, which could in future be solved by producing GWASs of individual symptoms, which would allow to better understand the relationship between phenotypes in these clusters.

## Conclusion

Navigome (phenviz.navigome.com) gathers visualizations of gene associations, expression, pathways, and genetic correlations for multiple human traits, based on GWAS results. The Navigome phenome map illustrates the correlation structure between phenotypes explained by common genetic variants, and can be divided into two broad regions: one for phenotypes related to greater longevity, height, education, and metabolites in HDL particles, and the other related to increased BMI, metabolic and mental disorders, coronary artery disease and lower longevity. Well-defined clusters could be identified on the map, which exhibit characteristic biological pathways. The map itself should be interpreted with caution, depending on the significance of the genetic correlations and map topology. Such a map could be used in the future with more traits, especially metabolic traits, to find biomarkers for known disorders.

## Methods

### Phenotype collection and quality control

We collected systematically 487 studies, some directly from websites of large consortia, some others designed by our group using UK Biobank data (the quality control procedure is described in Supplementary Text 1). These UK Biobank GWAS include: neuroticism, BMI, fat mass, lean mass, and physical activity (cf. following paragraphs). The GWAS from the Translational and Neuropsychiatric Genetics group (TNG) at King’s College London can be downloaded in Navigome (phenviz.navigome.com). All studies in Navigome contain signed summary statistics, which is a requirement to compute genetic correlations using LDSC regression as well as tissue-specific gene associations with S-PrediXcan. For secondary statistical analyses, we only used variants with known rs ids. We applied several filters to the genotype variants: (1) MAF ≥ 1%, (2) p-value between 0 and 1, (3) imputation info score ≥ 0.6 (when available), and (4) no unrealistic odds ratio (> 10,000). Effect alleles were identified systematically; it should be noted that confusion between A1 and A2 alleles is a recurrent problem when sharing GWAS summary statistics. When in doubt, the effect allele can be checked in dbGAP or genetic correlations can be computed to verify the direction of association with related phenotypes. If the effect allele (A1) and rs id were included but not the other allele (A2), we recovered A2 from 1000 Genomes phase III. Similarly, if genomic position was included as well as A1 and A2 but not the rsid, we recovered the rs id from 1000 Genomes phase III. The number of individuals per SNP was not provided in most studies, but was taken into account whenever available - otherwise, the total number of individuals or the total number of cases and controls (depending on the study design) was used.

### UK Biobank GWAS: physical activity

The GWAS on physical activity (Navigome code: PHYS01) uses imputed genotypes in 29,496 male and 36,758 female (N = 66,254) individuals in the UK Biobank. Covariates included age at recruitment, genotyping array, and 20 first principal components of the genotype matrix. Physical activity was measured over a period of 7 days with a wrist-worn accelerometer^9^. A wear-time adjusted 7-day average measure of activity was used, including only individuals meeting UK Biobank quality criteria: good wear-time, good calibration, calibration on own data, and no problem indicators. Analyses were performed on the intersection of this UK Biobank subset with those passing general genotyping quality control: in white British ancestry subset, used in the calculation of ancestry principal components, without excess relatives in the UK Biobank sample, no putative sex chromosome aneuploidy, and not outliers for heterozygosity and genotype missingness. General genotyping considerations, raw data quality control, and imputation procedure in the UK Biobank are described elsewhere^10^.

### UK Biobank GWAS: Body composition

We performed GWAS on BMI, body fat percentage, fat mass and lean mass using BGENIE v1.2 (https://jmarchini.org/bgenie) in 353,972 European participants from the UK Biobank. The corresponding study codes on Navigome are BODY08, BFPC04, FATM04, and LEAN06. Body composition was assessed using Tanita BC-418 MA scale (Tanita Corporation, Arlington Height, IL). 7,794,483 genotyped and imputed SNPs were included and insertion-deletion variants with a MAF of 1%. Pregnant participants or females after hysterectomy were excluded. Covariates included in the association analysis were assessment center, genotyping batch, smoking status, alcohol consumption, menopause, age, and Townsend Deprivation Index^11^. Population stratification was accounted for by including the first six principal components of the genotype matrix, from the European subsample.

### UK Biobank GWAS: Neuroticism

We performed sex-specific GWAS on neuroticism using the genotype data supplied by UK Biobank in males (N = 142,875) and in females (N =144,660; total N = 287,535. The corresponding Navigome codes are NEUR02B for both sexes, NEUR02F for females and NEUR02M for males. The neuroticism phenotype was calculated as the sum score of neuroticism questions at the baseline assessment^12^, corrected for age and sex-specific means and SDs from the UK population^13^. In a second analysis, individuals were excluded if they reported any psychiatric illness resulting in 83,413 males and 73,946 females (total N = 157,355). The corresponding Navigome codes are NEUR03B for both sexes, NEUR03F for females and NEUR03M for males. Sex-stratified linear regressions were performed in PLINK^14^ using eight principal components and genotyping batch as covariates and later meta-analyzed using METAL^15^.

### Map of genetic correlations

We computed genetic correlations between all phenotypes using LDSC v1.0.0^2^, using imputation info score ≥ 0.6 and otherwise default parameters. After computing all genetic correlations between the 487 studies using LDSC regression, we removed studies with more than 20% missing values and reached the final number of 465 studies. Missing values were imputed (median value imputation) before applying principal component analysis, which reduced the data dimensionality from 465 to 47 (47 components explaining 90% of the data variance). A two-dimensional map was generated from the complete correlation matrix using t-distributed stochastic neighbour embedding^8^ (t-SNE) implemented in the python package scikit-learn^16^, with default options: perplexity = 30, early exaggeration = 12, learning rate = 200, number of iterations = 1000. Other maps were also generated with different perplexity parameters (perplexity = {5, 10, 20, 30, 40, 50}) within the range recommended by Matten et al^8^, to see if clusters were preserved and make topological observations.

### Gene associations

We performed two types of gene-wise associations: (1) with MAGMA^6^, (2) with S-PrediXcan^7^. For a given phenotype, S-PrediXcan predicts changes in expression in different tissues, using expression quantitative trait loci (eQTL), which are loci in the genome linked to the variation in expression of mRNAs. MAGMA, on the other hand, provides a simple gene association test based on a combination of SNP association statistics^6^, which does not give information on gene expression.

Most GWAS results provide p-values for each tested variant (mainly SNPs) and a signed summary statistic such as the odds ratio or regression coefficient. To compute gene associations, MAGMA v1.06 only requires the SNP p-values, and computes gene associations using Brown’s method accounting for linkage disequilibrium using SNP genotype correlations derived from a reference population (here, 1000 Genomes Phase III)^17^.

S-PrediXcan, on the other hand, derives a Z-score or Wald statistic using (1) weights of SNPs in the prediction of gene expression derived from an expression training set (here, GTEx v7^18^ and Depression Genes and Networks cohort (DGN)^19^), (2) SNP variances and covariances derived from a reference population set (1000 Genomes Phase III^17^), (3) signed summary statics (not only the p-value). The sign of the Z-score indicates whether a gene could be upregulated or downregulated for a specific phenotype-tissue combination - for example, if calcium channels would be expected to be differentially expressed in schizophrenia cases versus controls.

### Pathway analysis

The set of 7267 pathways was constructed with MSigDB v6.1^20^ canonical pathways, GO ontology, and a small set of pathways from the literature (significant pathways are referenced on phenviz.navigome.com). Pathway analysis was performed using MAGMA v.1.06 competitive analysis^6^. This type of analysis, based on gene-wise associations obtained using Brown’s method, tests whether genes within a specific pathway are more associated with the phenotype than genes outside of the pathway, whilst correcting for confounders such as gene size, gene density and minor allele count.

### Significance thresholds for multiple testing

Considering a rounded number of 20,000 tested protein-coding genes, a fixed Bonferroni threshold can adopted for MAGMA gene testing (*p-value* ≤ 0.05/20000 = 2.5E-6). For PrediXcan associations, since the gene association test is run for several tissues, a more stringent threshold should be used, accounting for ~258,500 gene-tissue pairs (*p-value* ≤ 1.93E-7). However, a p-value ≤ 2.5E-6 threshold can be used to compare MAGMA and S-PrediXcan results. For pathways, the threshold is p ≤ 1E-5 ~ 0.05/*N_indep_*, where *N_indep_* = 5254 is the number of independent tests. We estimated *N_indep_* by (1) computing the 7267×7267 Tanimoto similarity matrix of pathways (2) running principal component analyses to find the number of principal components explaining 99.5% of the data variance, which is the final *N_indep_* estimate.

## Acknowledgments

All GWAS summary statistics used in this study are referenced in the Navigome web interface (phenviz.navigome.com) and Supplementary Table. GWAS summary statistics from our group at King’s College London are accessible for download on the Navigome website (phenviz.navigome.com/downloads). HG and GB acknowledge funding from the US National Institute of Mental Health (PGC3: U01 MH109528). We gratefully acknowledge capital and computing equipment funding from the Maudsley Charity (Grant Reference 980) and Guy’s and St Thomas’s Charity (Grant Reference STR130505).

### Conflict of interest

GB reports consultancy and speaker fees from Eli Lilly and Illumina and grant funding from Eli Lilly.

## References

1. Sullivan, P. F. et al. Psychiatric Genomics: An Update and an Agenda. Am. J. Psychiatry 175, 15–27 (2018).

2. Brendan Bulik-Sullivan, H. F. LD Score Regression (LDSC). (Broad Institute of MIT and Harvard / MIT Department of Mathematics, 2015).

3. Bulik-Sullivan, B. et al. An atlas of genetic correlations across human diseases and traits. Nat. Genet. 47, 1236–1241 (2015).

4. Zheng, J. et al. LD Hub: a centralized database and web interface to perform LD score regression that maximizes the potential of summary level GWAS data for SNP heritability and genetic correlation analysis. Bioinformatics 33, 272–279 (2017).

5. Lee, P. H., O’Dushlaine, C., Thomas, B. & Purcell, S. M. INRICH: interval-based enrichment analysis for genome-wide association studies. Bioinformatics 28, 1797–1799 (2012).

6. de Leeuw, C. A., Mooij, J. M., Heskes, T. & Posthuma, D. MAGMA: generalized gene-set analysis of GWAS data. PLoS Comput. Biol. 11, e1004219 (2015).

7. Barbeira, A. N. et al. Exploring the phenotypic consequences of tissue specific gene expression variation inferred from GWAS summary statistics. Nat. Commun. 9, 1825 (2018).

8. Maaten, L. Visualizing High-Dimensional Data Using t-SNE. J. Mach. Learn. Res. 9, 2579–2605 (2008).

9. Doherty, A. et al. Large Scale Population Assessment of Physical Activity Using Wrist Worn Accelerometers: The UK Biobank Study. PLoS One 12, e0169649 (2017).

10. Bycroft, C. et al. Genome-wide genetic data on ~500,000 UK Biobank participants. bioRxiv 166298 (2017). doi:10.1101/166298

11. Townsend, P. Deprivation*. J. Soc. Policy 16, 125–146 (1987).

12. Smith, D. J. et al. Prevalence and characteristics of probable major depression and bipolar disorder within UK biobank: cross-sectional study of 172,751 participants. PLoS One 8, e75362 (2013).

13. Eysenck, S. B. G., Eysenck, H. J. & Barrett, P. A revised version of the psychoticism scale. Pers. Individ. Dif. 6, 21–29 (1985).

14. Purcell, S. et al. PLINK: A Tool Set for Whole-Genome Association and Population-Based Linkage Analyses. in Am J Hum Genet 81, 559–575 (2007).

15. Willer, C. J., Li, Y. & Abecasis, G. R. METAL: fast and efficient meta-analysis of genomewide association scans. Bioinformatics 26, 2190–2191 (2010).

16. Buitinck, L. et al. API design for machine learning software: experiences from the scikit-learn project. arXiv [cs.LG] (2013).

17. 1000 Genomes Project Consortium et al. A global reference for human genetic variation. Nature 526, 68–74 (2015).

18. GTEx Consortium. The Genotype-Tissue Expression (GTEx) project. Nat. Genet. 45, 580–585 (2013).

19. Battle, A. et al. Characterizing the genetic basis of transcriptome diversity through RNA-sequencing of 922 individuals. Genome Res. 24, 14–24 (2014).

20. Liberzon, A. et al. The Molecular Signatures Database (MSigDB) hallmark gene set collection. Cell Syst 1, 417–425 (2015).

